# Genomic prediction of celiac disease targeting HLA-positive individuals

**DOI:** 10.1101/017608

**Authors:** Gad Abraham, Alexia Rohmer, Jason A. Tye-Din, Michael Inouye

**Affiliations:** Centre for Systems Genomics, School of BioSciences, The University of Melbourne, Parkville 3010, Victoria, Australia; Medical Systems Biology, Department of Pathology and Department of Microbiology & Immunology, The University of Melbourne, Parkville 3010, Victoria, Australia; Faculty of Life Science, University of Strasbourg, 67084 Strasbourg CEDEX, France; The Walter and Eliza Hall Institute of Medical Research, 1G Royal Parade, Parkville 3052, Victoria, Australia; Department of Medical Biology, The University of Melbourne, Parkville 3010, Victoria, Australia; Department of Gastroenterology, The Royal Melbourne Hospital, Grattan St., Parkville 3050, Victoria, Australia; Murdoch Children’s Research Institute, Flemington Road, Parkville, Victoria 3050, Australia

## Abstract

**Background:** Genomic prediction aims to leverage genome-wide genetic data towards better disease diagnostics and risk scores. We have previously published a genomic risk score (GRS) for celiac disease (CD), a common and highly heritable autoimmune disease, which differentiates between CD cases and population-based controls at a clinically-relevant predictive level, improving upon other gene-based approaches. HLA risk haplotypes, particularly HLA-DQ2.5, are necessary but not sufficient for CD, with at least one HLA risk haplotype present in up to half of most Caucasian populations. Here, we assess a genomic prediction strategy that specifically targets this common genetic susceptibility subtype, utilizing a supervised learning procedure for CD that leverages known HLA-DQ2.5 risk.

**Methods:** Using L1/L2-regularized support-vector machines trained on large European case-control datasets, we constructed novel CD GRSs specific to individuals with HLA-DQ2.5 risk haplotypes (GRS-DQ2.5) and compared them with the predictive power of the existing CD GRS (GRS14) as well as two haplotype-based approaches, externally validating the results in a North American case-control study.

**Results:** Consistent with previous observations, both the existing GRS14 and the GRS-DQ2.5 had better predictive performance than the HLA haplotype approaches. GRS-DQ2.5 models, based on directly genotyped or imputed markers, achieved similar levels of predictive performance (AUC = 0.718—0.73), which were substantially higher than those obtained from the DQ2.5 zygosity alone (AUC = 0.558), the HLA risk haplotype method (AUC = 0.634), or the generic GRS14 (AUC = 0.679). In a screening model of at-risk individuals, the GRS-DQ2.5 lowered the number of unnecessary follow-up tests for CD across most sensitivity levels. Relative to a baseline implicating all DQ2.5-positive individuals for follow-up, the GRS-DQ2.5 resulted in a net saving of 2.2 unnecessary follow-up tests for each justified test while still capturing 90% of DQ2.5-positive CD cases.

**Conclusions:** Genomic risk scores for CD that target genetically at-risk sub-groups improve predictive performance beyond traditional approaches and may represent a useful strategy for prioritizing individuals at increase risk of disease, thus potentially reducing unnecessary follow-up diagnostic tests.

## Background

Genome-wide association studies (GWAS) have identified large numbers of genetic loci associated with complex human disease, particularly for many autoimmune diseases, where disease susceptibility is typically strongly linked to the Human Leukocyte Antigen (HLA) region as well as loci outside HLA [1-6]. The strong disease associations of specific single nucleotide polymorphisms (SNP) have enabled genomic-based prediction models to be developed with substantial predictive power [7-12]. As new light continues to be shed on the fundamental role of these genetic links in disease pathogenesis, it is becoming increasingly likely that genomic-based tools to predict disease development and risk, as well as prognosis and clinical course, may be harnessed and applied with direct relevance to patient care.

Despite these advances, suitable clinical tools that quantify genomic risk for complex disease remain largely unrealized. This is largely due to a lack of understanding regarding how such tools might be utilized in clinical settings, hampered by the complexity of integrating such data into a risk model that often incorporates many other variables, such as clinical information and laboratory investigations. The role of genomic risk prediction in existing risk models and diagnostic pathways is still being determined, including whether genomic prediction is optimal as a complement or a replacement for existing assays. For autoimmune diseases in particular, genomic prediction needs to demonstrate improved risk stratification beyond known HLA risk haplotypes and have sufficient predictive power, particularly given the low disease prevalence in the general population (typically 1% or less). Here, we have sought to identify specific scenarios where there is a major clinical need to improve risk prediction in a highly heritable autoimmune disease, celiac disease (CD). We first show that genomic risk prediction methods have clear advantages over existing approaches for CD risk prediction, and next assess the clinical implications of genomic risk prediction for disease management.

Celiac disease (CD) is a common systemic autoimmune disease caused by dietary gluten in genetically susceptible individuals [13-15]. CD affects ∼1% of the Western world and is strongly heritable (∼80% on the liability scale) [16]. The major genetic association is in the MHC locus, with specific HLA haplotypes present in almost all (∼99.6%) cases: HLA-DQ2.5 (*DQA1*05* / *DQB1*02*) in ∼88%, HLA-DQ2.2 (*DQA1*02* / *DQB1*02*) in ∼4%, and/or HLA-DQ8 (*DQA1*03 /DQB1*03:02*) in ∼6% [17]. This HLA association underpins the crucial pathogenic role of CD4+ T cells targeting a restricted repertoire of immunogenic gluten peptides [18]. Recent GWAS in CD implicate at least 41 other non-HLA loci with more modest contributions to risk [3, 4, 6, 19-22]. These regions are all linked to aspects of immune system function, and are likely to impact on CD susceptibility or clinical behavior.

Current diagnosis of CD relies on the presence of CD-specific autoantibodies and a confirmatory small-bowel biopsy to demonstrate the characteristic intestinal villous atrophy [13]. Importantly, while both methods are useful for detecting current CD, they do not provide predictive information on the future risk of developing CD in a person without active disease. Early detection of CD is a clinical priority in order to reduce long-term risk of disease complications, especially for individuals at higher-risk of CD, such as those who are a 1^st^-degree family members of an affected individual or those who have a related autoimmune disease. However, the follow-up care of people without active CD, particularly the question of whether and how often to perform follow-up testing, remains unresolved. This challenge stems from the desire for early disease detection conflicting with the need for minimizing repeated testing that is inconvenient, costly, anxiety-inducing, and entirely unnecessary for those individuals who will never develop disease.

Since the development of CD depends so strongly on several HLA risk haplotypes, HLA typing is able to achieve close to 100% sensitivity in detecting at-risk individuals while also excluding those at very low risk. As a result, HLA typing for CD has been widely embraced in clinical practice [23]. Indeed, recent consensus guidelines recommend HLA typing as a 1^st^ line investigation in asymptomatic children at-risk of CD (for instance, if they have a family history of CD) [13]. Further, combining HLA typing with CD-serology may also provide a more cost-effective diagnostic approach in some situations such as population screening by identifying false-positive serology in non-genetically susceptible individuals, thus reducing the number of unnecessary and expensive confirmatory small bowel biopsies [24].

The presence of at least one of the CD-related HLA haplotypes, while necessary for the development of disease, carries little predictive value for eventual CD development and therefore has no role as a sole diagnostic for CD. The CD-related genotypes are highly prevalent in the community, with 30—40% of Europeans and up to 56% Australians carrying at least one [24]. Despite the fact that different HLA haplotypes impart varying levels of risk for CD, specifically HLA-DQ2.5 > DQ8 > DQ2.2, these differences in relative risk have not translated meaningfully into clinical practice. Thus HLA typing results are typically interpreted as simply “susceptibility present” or “susceptibility absent” regardless of the particular HLA haplotype detected, and therefore used primarily to exclude CD but not to detect high-risk individuals nor to predict future risk of disease [13].

Notwithstanding the limited clinical role of HLA typing, there has been renewed interest in the role of specific HLA alleles in the development on CD, with the risk of CD being far higher in children who are HLA-DQ2.5 homozygous (or have two copies of *DQB1*02*) than among those who are HLA-DQ2.5 heterozygous or are positive for HLA-DQ8 [25, 26]. A gene-dose effect has been reported, with HLA-DQ2.5 homozygosity associated in some studies with a more severe clinical presentation of CD, refractory disease, and a slower rate of intestinal healing on treatment [27, 28]. As a result, HLA-DQ2.5 zygosity status is a major variable in a recently proposed CD prognostic modeling tool [29]. Collectively, these studies highlight the important role of the HLA risk haplotypes, especially HLA-DQ2.5, in CD development, prognosis, and clinical behavior, but also highlight the current limitations in the clinical utility of HLA typing. There is a major need to develop tools that are more informative than HLA typing particular for those who have already been shown to possess at least one HLA-DQ2.5 allele (DQ2.5+).

We have recently performed a proof-of-principle study demonstrating that genomic data, derived from multiple case/control GWAS datasets, can be used to improve upon current genetic testing based on HLA typing [7]. A CD genomic risk score (GRS) based on genome-wide single nucleotide polymorphisms (SNPs), denoted here the “GRS14” [8], was induced by supervised learning models trained on a British case-control GWAS study [30, 31]. We have established the robustness of GRS14 to discriminate CD patients from population-based controls in UK, Dutch, Finnish, and Italian studies [3, 4, 32], achieving predictive performance (Area Under the receiver-operating characteristic Curve (AUC)) substantially higher than other methods which attribute risk based on 57 non-HLA SNPs together with HLA alleles [33].

A recent large-scale CD Immunochip GWAS [34] has shown that statistical imputation of non-SNP genetic variants via SNP2HLA [35], including HLA alleles and amino-acid polymorphisms in HLA genes, enabled fine-mapping of CD-associated variation to a greater resolution than that afforded by SNPs alone, and subsequently substantially increased the explained heritability of CD. However, the implications for CD risk prediction had not been assessed.

Here, we first externally validate the predictive power of the existing GRS14 and novel variants thereof in a North American CD case-control study [22, 36], comparing their performance to a previous HLA haplotype risk approach. Next, we focus on the DQ2.5+ subset of individuals, and develop HLA-DQ2.5-specific genomic risk scores, one based on SNPs and others based on SNPs together with SNP2HLA-imputed markers [35]. Finally, we assess the HLA-DQ2.5-specific genomic risk scores in screening scenarios to determine the number of unnecessary follow-up tests saved relative to other approaches.

## Methods

### Genotype and phenotype data

We obtained the North American dataset (cases and controls) from NCBI dbGaP (accession phs000274.v1.p1). Samples were genotyped on the Illumina 660W Quad v1A platform, assaying 2246 individuals in total (1716 cases, 530 controls, 723 male, 1523 female). Individuals were considered to have confirmed CD based on either (i) characteristic findings on small bowel biopsy according to ESPGHAN criteria, (ii) biopsy-proven dermatitis herpetiformis, or (iii) positive celiac serology panel (transglutaminase (tTG) and endomysial (EMA) antibodies). Controls were originally from the Illumina iControl database, matched by the original study authors for age, sex, and ethnicity [36]. To minimize the possibility of artificially inflating the apparent predictive ability of the models [37], we performed several stages of quality control on the genotype data using plink 1.9 (https://www.cog-genomics.org/plink2) [38, 39]: removing non-autosomal SNPs, filtering SNPs by minor allele frequency (MAF) <1%, missingness >10%, deviation from Hardy-Weinberg equilibrium in controls *P* <5×10^−6^, and filtering of individuals with missingness >10%. Next we removed 2473 SNPs with case/control SNP differential missingness *P* <10^−3^. We iteratively used principal component analysis (PCA), implemented in flashpca 1.2 [40], to identify outlier individuals, defined here as individuals with PC coordinates more than 3 standard deviations from the median of each of PC 1—50. After removing those individuals, PCA was repeated to verify the results. We used two iterations of this procedure, resulting in 1697 individuals remaining. Finally, we removed one of two individuals with identity-by-descent 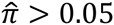 (one individual was removed). The final QCd dataset consisted of 1696 individuals (1259 cases, 437 controls, 546 males and 1150 females) over 518,770 autosomal SNPs. The available clinical characteristics of the post-QC data are shown in Table 1.

**Table 1:**
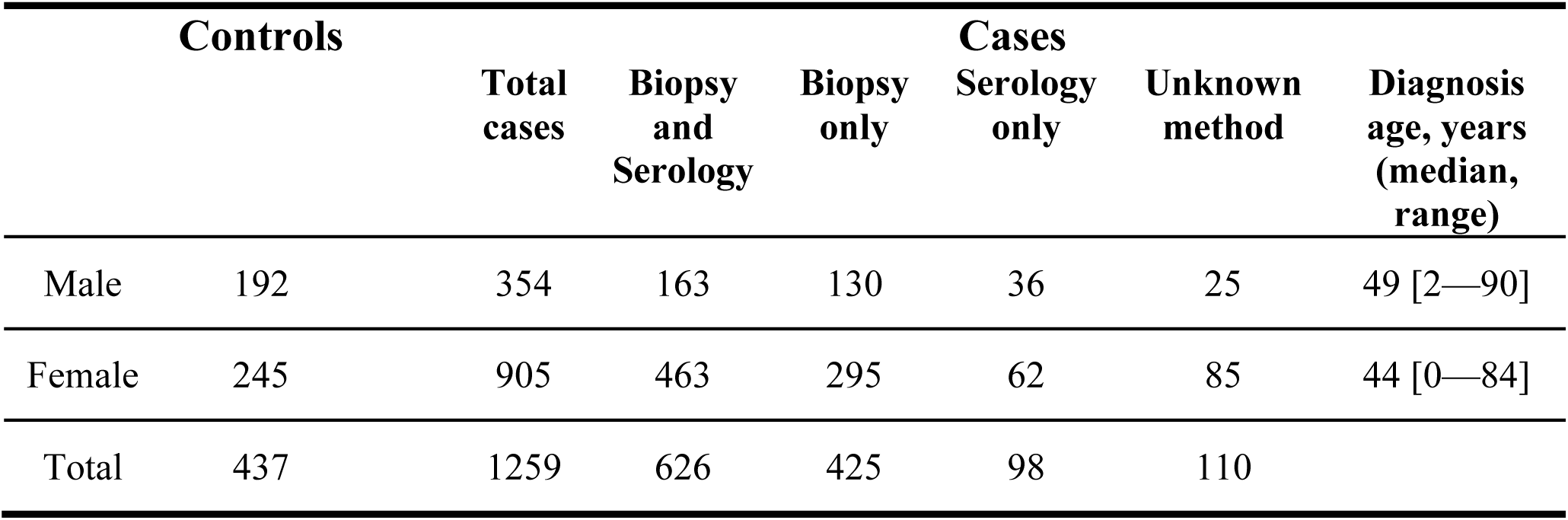
The North American NIDDK-CIDR dataset clinical characteristics (post QC).

The genotype data for the UK (n=6785), Finnish (n=2476), Dutch (n=1649), and Italian (n=1040) cohorts have been previously described [3, 4, 8, 32]; these datasets were genotyped on the Illumina 670-QuadCustom-v1, 610-Quad, and 1.2M-DuoCustom-v1 genome-wide SNP arrays. Each cohort underwent additional separate QC: removing SNPs with MAF <1%, Hardy-Weinberg deviation from equilibrium in controls *P* < 5×10^−6^, missingness >10%, or differential case/control missingness *P* < 10^−3^, and removing samples with missingness >10% or IBD 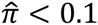, before being combined into a single dataset consisting of n=11,912 samples and 500,821 SNPs. The Immunochip dataset (n=16,002) was assayed on a custom Illumina fine-mapping array (Immunochip) comprising 115,746 SNPs after QC, as described [3, 8] (with SNPs further filtered by MAF >0.5%), of which 17,848 were common to both the Immunochip and GWA arrays. Since some individuals were genotyped both in the UK GWA dataset and in the Immunochip dataset, when combining the GWA and Immunochip datasets we included only individuals with pairwise identity-by-descent (plink IBD) 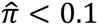, resulting in n=19,715 individuals in total.

In order to check whether the putative European descent of the North American individuals was indeed the case, we also combined an LD-thinned (plink – indep-pairwise) version of the North American data with an LD-thinned version of the UK, Finnish, Dutch, and Italian GWA data and performed PCA on the combined data (**Additional File 1**, Supplementary Figure 1). Finally, we computed the fixation index F_st_ [41] (plink-fst) on a combined dataset (European + North American) consisting of the 224 of the 228 SNPs in the GRS14, with the European and North American samples as two clusters.

### Ethics statement

All participants gave informed consent and the study protocols were approved by the relevant institutional or national ethics committees. Details for the ethics protocols for the European GWA and Immunochip datasets are given in [3, 4]. The North American NIDDK-CIDR dataset was obtained from the NCBI Database of Genotypes and Phenotypes (dbGaP), accession phs000274.v1.p1, following their respective access protocols.

### Imputation of HLA alleles and other markers

While SNP arrays only produce unphased alleles, statistical methods can exploit patterns of linkage disequilibrium to impute haplotypes (phased alleles) and other forms of genetic variation, based on a reference panel where both the SNPs and the imputed variation are known (a training dataset). This procedure is conceptually the same as used when imputing, for example, millions of 1000Genome genotypes based on a smaller set of hundreds of thousands of assayed genotypes.

As the majority of CD-related variation is found within the HLA, we imputed HLA variation, employing two complementary methods. First, we imputed *HLA-DQA1* and *HLA-DQB1* alleles from the SNPs using the R package HIBAG 1.2.3 [42], using the European hg18 HLA4 reference dataset. Based on the imputed HLA alleles, we inferred each individual’s heterodimer type as one of DQ2.5 heterozygous, DQ2.5 homozygous, DQ2.2, or DQ8, according to the mapping in ref [8]. Following [33], the HLA risk score was assigned as low for individuals that did not have any of the CD risk heterodimers (DQ2.2, DQ8, DQ2.5-heterozygous, and DQ2.5-homozygous). High risk was assigned to individuals with DQ2.5-homozygous or those with both DQ2.5-heterozygous and DQ2.2. Medium risk was assigned to all other remaining individuals. The HLA risk profiles were coded as 0 for low, 1 for medium, and 2 for high risk. We did not examine the 57 non-HLA SNPs used in [33] as these are only present on Immunochip arrays and not on the genome-wide arrays, and were not well tagged by the existing SNPs on the genome-wide arrays.

Another imputation approach, SNP2HLA [35], has proven to be particularly useful for fine-mapping of the HLA region with regards to association with several autoimmune diseases, including CD [34], and for explaining more of the heritability of disease. SNP2HLA employs the assayed SNPs, together with a reference panel, to impute HLA SNPs (if they were not already on the genotyping array), HLA alleles (*HLA-A*, *HLA-B*, *HLA-C*, *HLA-DPA1*, *HLA-DPB1*, *HLA-DQA1*, *HLA-DQB1*, and *HLA-DRB1*), and known amino acid polymorphisms within those HLA genes. These imputed variants can then be analyzed in the same way as assayed genotypes.

Hence, in addition to using HIBAG, we employed SNP2HLA v1.0.2 [35] to impute 8961 HLA SNPs, eight HLA alleles, and amino acid polymorphisms, based on the T1DGC reference panel, in the European GWA, Immunochip, and North American dataset. The non-SNP imputed markers were coded as present/absent. Quality control for the combined SNP + imputed marker data included (i) removal of imputed SNPs that were already assayed on the array; (ii) within each dataset (UK2, NL, Finn, IT), removal of SNPs/markers with MAF <1%, missingness >10%, deviation from Hardy-Weinberg equilibrium (HWE) in controls *P* <5×10^−6^, differential case/control missingness *P* <10^−3^, and removal of individuals with >10% missingness; (iii) removal of SNPs/markers that were not present in the four European datasets (UK2, NL, Finn, IT); (iv) removal of SNPs/markers with differential case/control missingness *P* <10^−3^ across the combined data; and (v) removal of SNPs/markers not on the North American imputed dataset. For the Immunochip data, QC included (i) removal of imputed SNPs already assayed; (ii) SNP/marker filtering by MAF <0.5%, missingness >10%, deviation from HWE in controls *P* <10^−3^, and differential case/control missingness *P* <10^−3^, and removal of individuals with missingness >10%. We verified that the imputed markers included in genomic risk scores had high imputation accuracy (*r*^2^ >0.8) in the training data. The final SNPs + imputed marker European data consisted of 507,321 markers (500,821 assayed SNPs and 6500 imputed markers) over 11,912 individuals (5552 of which were DQ2.5+); after removal of individual assayed in the UK GWA data, the Immunochip dataset had 7803 individuals, of which 4732 were DQ2.5+, with 24,555 SNPs/markers (∼17,800 assayed SNPs and ∼6700 imputed markers) common with the other datasets. For the DQ2.5-specific GRS (GRS-DQ2.5), only SNPs present in the North American dataset were used in cross-validation, so that all SNPs present in the model could be used to determine the score in external validation. SNP2HLA had ∼100% concordance with HIBAG’s DQ2.5+ classification.

### Validation of the published risk score

We used the previously published GRS14 risk score (comprising 228 SNPs, available at http://dx.doi.org/10.6084/m9.figshare.154193), which is given in terms of rs IDs, reference alleles, and a weight, to produce a per-individual score (using plink-score). The final score for each individual is the sum of the minor allele dosages of each SNP, weighted by the published weights. Four SNPs in this score were not found in the North American post-QC genotype data and were excluded; these four SNPs had relatively low weight (ranked 59th or lower, out of 228) and thus their absence is unlikely to have substantially affected the predictive power of the final model.

### Cross-validation and novel genomic risk scores

We used the tool SparSNP [30], which fits L1/L2-penalized support-vector machine (SVM) models to SNP data, in 10×10-fold cross-validation on the European GWA dataset. Briefly, these models are additive in the minor allele dosage {0, 1, 2}, and take into account all SNPs (or other markers) in the data, however, only a proportion of the SNPs/markers receive a non-zero weight, tuned by the L1 penalty (higher penalties lead to fewer SNPs/markers with non-zero weight), together with an L2 (ridge) penalty varying from 10^−6^ to 10^3^. The optimal penalties were determined via cross-validated AUC. We have previously shown that such penalized models produce superior predictive ability for CD compared with several widely-used alternatives [12]. We evaluated a range of L1/L2-penalized models over a grid of penalties, with the optimal model selected by the best average AUC. For cross-validation, the reported AUC is a LOESS-smoothed average over the 10×10 = 100 test sets. For independent validation, we derived a final consensus model consisting of the SNPs selected in >60% of the replications, with corresponding weights being the average weights over the replications. The consensus model was taken and tested without further modification on the North American dataset. Improvement in case/control discrimination (AUC) was tested using Harrell’s two-sided test for paired concordance (rcorrp.cens in R package Hmisc) [43], and 95% confidence intervals for the AUC were computed using DeLong’s method (R package pROC) [44].

### The ratio of non-CDs incorrectly implicated per CD correctly implicated

We calculated the ratio *r* of non-CD individuals incorrectly implicated per CD case correctly implicated as r = (1 - PPV)/PPV, where *PPV* is the positive predictive value, calculated as

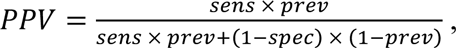

where *sens*, *spec*, and *prev* are the sensitivity, specificity, and prevalence (as a proportion of the population under consideration), respectively, computed for each possible risk cutoff (forming the ROC curve). This ratio is also the reciprocal of the post-test odds of disease, that is, 1 / (likelihood-ratio × pre-test-odds). In estimating the PPV for the DQ2.5+ subgroup we assumed a prevalence of 10%, equivalent to the prevalence of individuals with high background risk of CD, such as family history of the disease.

To evaluate the difference in the ratios between two risk scores, we used a stratified bootstrap procedure in the test data, whereby *B*=10,000 replications were drawn (sampled with replacement), the rank statistics were estimated within each replicate for each risk score separately, and the average over the differences in the ratios was reported as the final bootstrap estimate, with 95% approximate confidence intervals for the difference derived using the 0.025 and 0.975 quantiles of the bootstrapped differences.

## Results

### Independent validation of genomic risk scores for CD

We applied the previously published GRS14 to the North American dataset and evaluated its predictive power using receiver-operating characteristic (ROC) curves (Figure 1). The GRS14 model achieved AUC = 0.831 (95% CI 0.808—0.854), indicating that the majority of the predictive power of the GRS14 model, previously estimated at AUC = 0.86—0.9 [8] on the Italian, Dutch, and Finnish datasets, was maintained in the North American dataset. For comparison, we also trained MultiBLUP [11] on the same European data and tested it on the North American dataset with identical results (AUC = 0.831, 95% CI 0.808—0.85; **Additional File 1, Supplementary Figure 2),** and in addition employed the same L1/L2-regularized SVMs in cross-validation within the North American data, yielding similar results (maximum average cross-validated AUC = 0.823, **Additional File 1**, Supplementary Figure 3). Further, there were no substantial differences in the genomic scores between CD cases diagnoses with different diagnosis methods (**Additional File 1, Supplementary Results** and Supplementary Figure 4).

**Figure 1:**
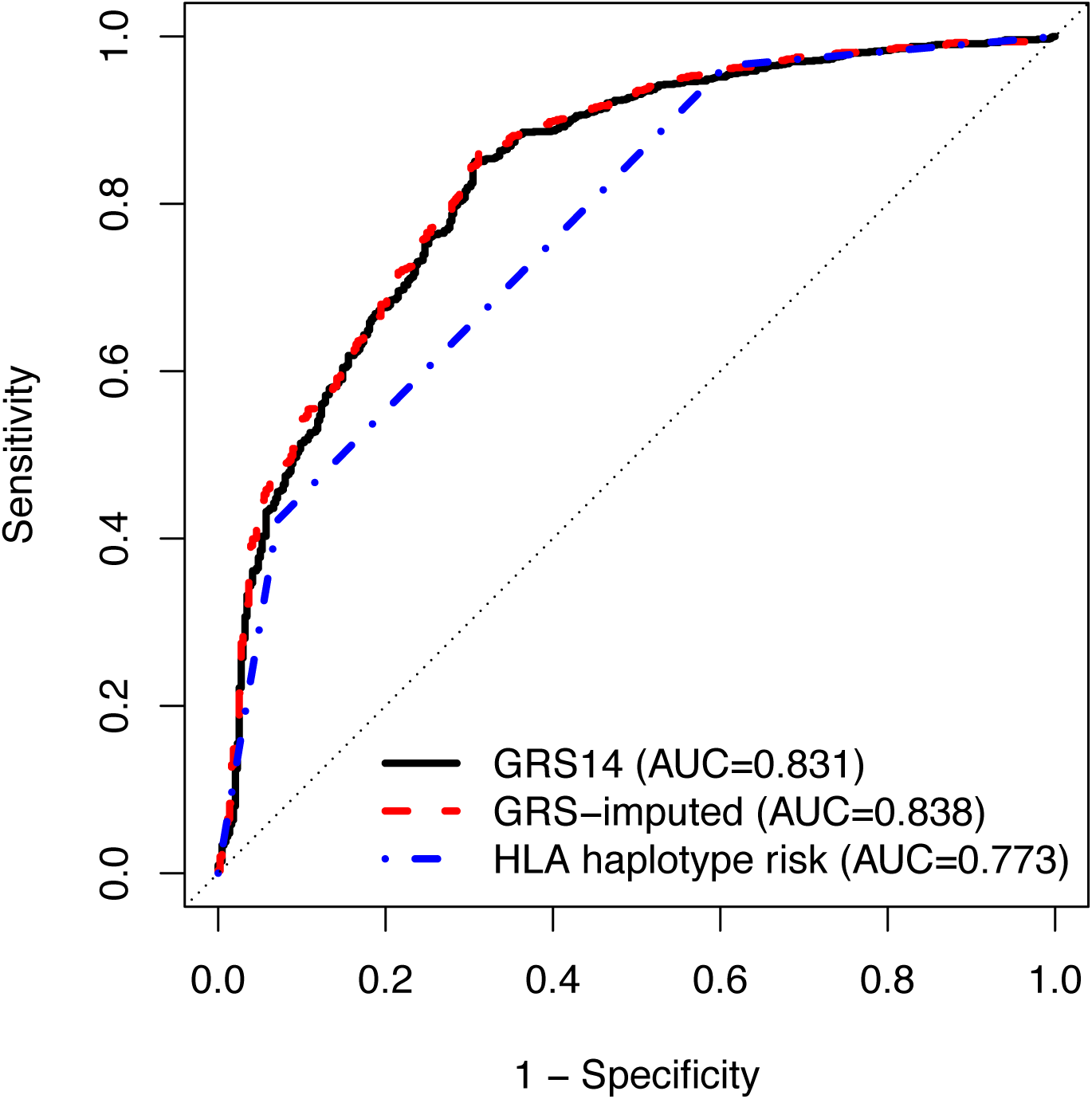
ROC curves for classifying all CD cases and controls using different predictors in the North American dataset. **Legend:** *GRS14*: the published GRS (trained on the UK2 dataset); *GRS-imputed:* a GRS trained on all European GWA datasets (UK, Dutch, Finnish, Italian), consisting of SNPs and SNP2HLA imputed markers; *HLA haplotype risk:* a 3-level risk score based on the HLA haplotype status.

In comparison to the GRS14, the 3-level haplotype risk method had substantially lower predictive power, with AUC = 0.773 (95% CI 0.751—0.795). The GRS model trained on the European GWA SNPs together with SNP2HLA imputed markers (“GRS-imputed”), improved the AUC over GRS14 by +0.007 (AUC = 0.838, CI 0.816—0.860, *P* <10^−6^ value for paired concordance test against the GRS14) (Figure 1). However, training a similar GRS-imputed model on the combined Immunochip + GWA dataset resulted in reduced performance relative to the GWA-only model (AUC = 0.835, 0.813—0.858, *P <*10^−6^ against the GRS-imputed model trained on the GWA data only), despite the larger sample size (**Additional File 1**, Supplementary Figure 2).

### Celiac disease risk prediction within the HLA-DQ2.5+ subgroup

While risk scores that discriminate CD cases from controls in the general population are useful, a more pressing clinical question is whether discrimination is possible within the HLA-DQ2.5+ subgroup of individuals, who are at the highest risk for CD amongst all HLA+ individuals. It is estimated that ∼90% of HLA+ individuals are DQ2.5+, with those that are DQ2.5-homozygous being at greater risk for CD than those that are DQ2.5-heterozygous [26, 45-47].

Restricting our analysis to DQ2.5+ individuals, we trained two new GRSs, one using a sparse linear model of SNPs only (GRS-DQ2.5), and another built similarly to GRS-DQ2.5 but also utilizing markers imputed by SNP2HLA (GRS-DQ2.5-imputed). These new GRS’s were then compared to three other predictive models:

1. the imputed DQ2.5 zygosity status for each individual (DQ2.5-zygosity),
2. the 3-level HLA haplotype risk score (HLA-haplotype-risk),
3. and the published GRS14.

The GRS-DQ2.5 and GRS-DQ2.5-imputed models were evaluated using the average AUC over 10×10-fold cross-validation on the DQ2.5+ European GWA samples. For GRS-DQ2.5, the maximum AUC achieved was 0.727 at 2513 SNPs with non-zero weight together with an L2 penalty of 1. For the GRS-DQ2.5-imputed, the best AUC was 0.74 at 3317 non-zero weight SNPs/markers using an L2 penalty of 1 as well (Figure 2) (for the results of GRS-DQ2.5 using other L2 penalties see **Additional File 1**, Supplementary Figure 5).

**Figure 2:**
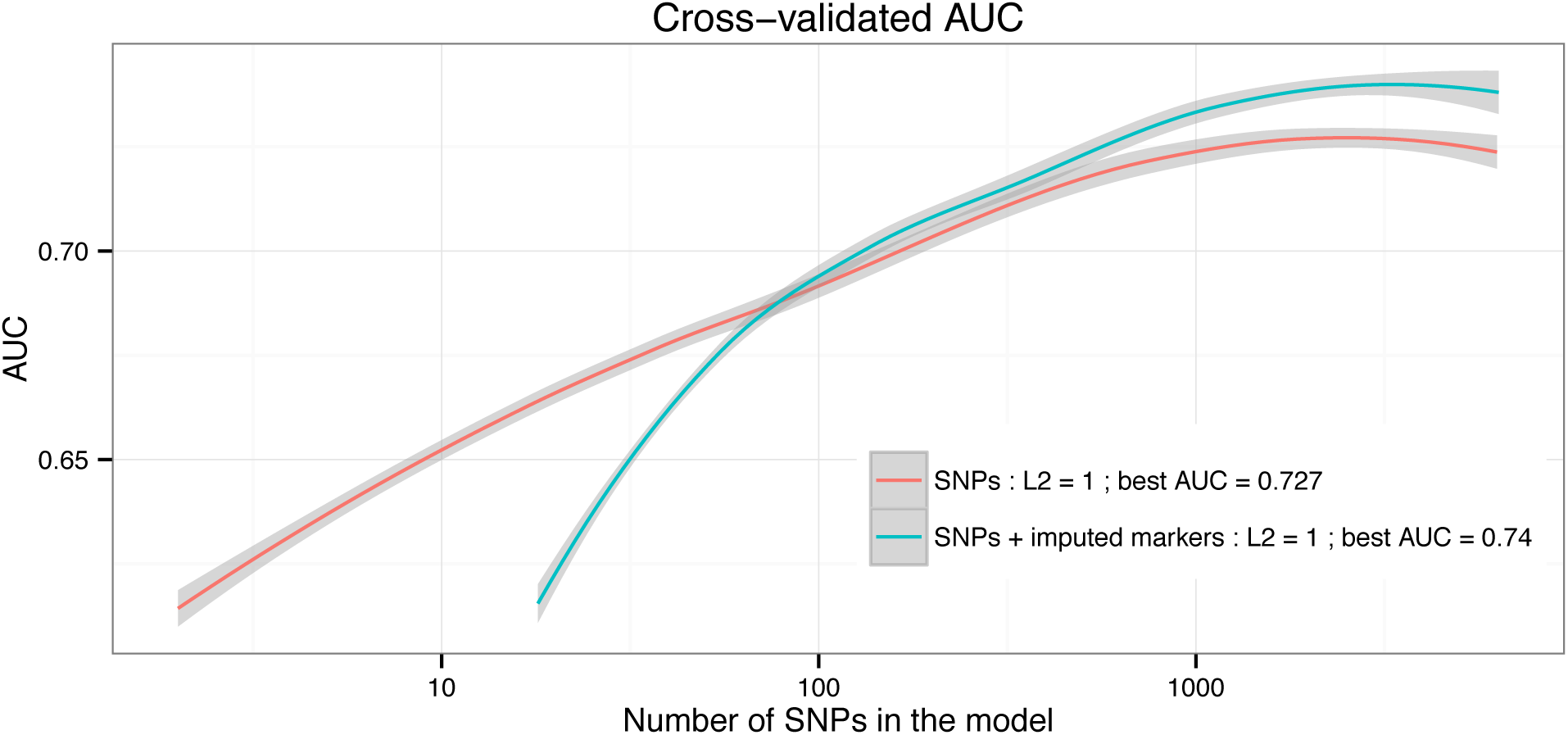
Cross-validation results for the GRS-DQ2.5 and GRS-DQ2.5-imputed models in the European GWA data. **Legend:** 10×10 cross-validated AUC (LOESS-smoothed) for the novel GRS-DQ2.5 model trained on the DQ2.5+ subset of the European GWA data (n=5552), as a function of the number of SNPs assigned a non-zero weight in the model. Maximum AUC was 0.727 achieved at 2513 SNPs with non-zero weight when considering only SNPs (GRS-DQ2.5) and AUC of 0.74 at 3317 SNPs/markers when using SNPs and SNP2HLA-imputed markers (GRS-DQ2.5-imputed).

To externally validate these models, we utilized the North American DQ2.5+ individuals (n=1237, 1094 cases and 143 controls) (Figure 3a). The highest performance was observed for GRS-DQ2.5-imputed, achieving an AUC of 0.73 (95% CI 0.687—0.772), followed by GRS-DQ2.5 with AUC = 0.718 (95% CI 0.676—0.761) and the GRS14 with AUC = 0.669 (95% CI 0.625—0.713). The HLA-haplotype-risk and DQ2.5-zygosity models achieved AUCs of 0.634 (95% CI 0.597—0.671) and 0.558 (95% CI 0.534—0.582), respectively. Training similar models on a combined Immunochip and GWA dataset resulted in lower externally validated AUC of 0.707 (**Additional File 1, Supplementary Results**).

**Figure 3:**
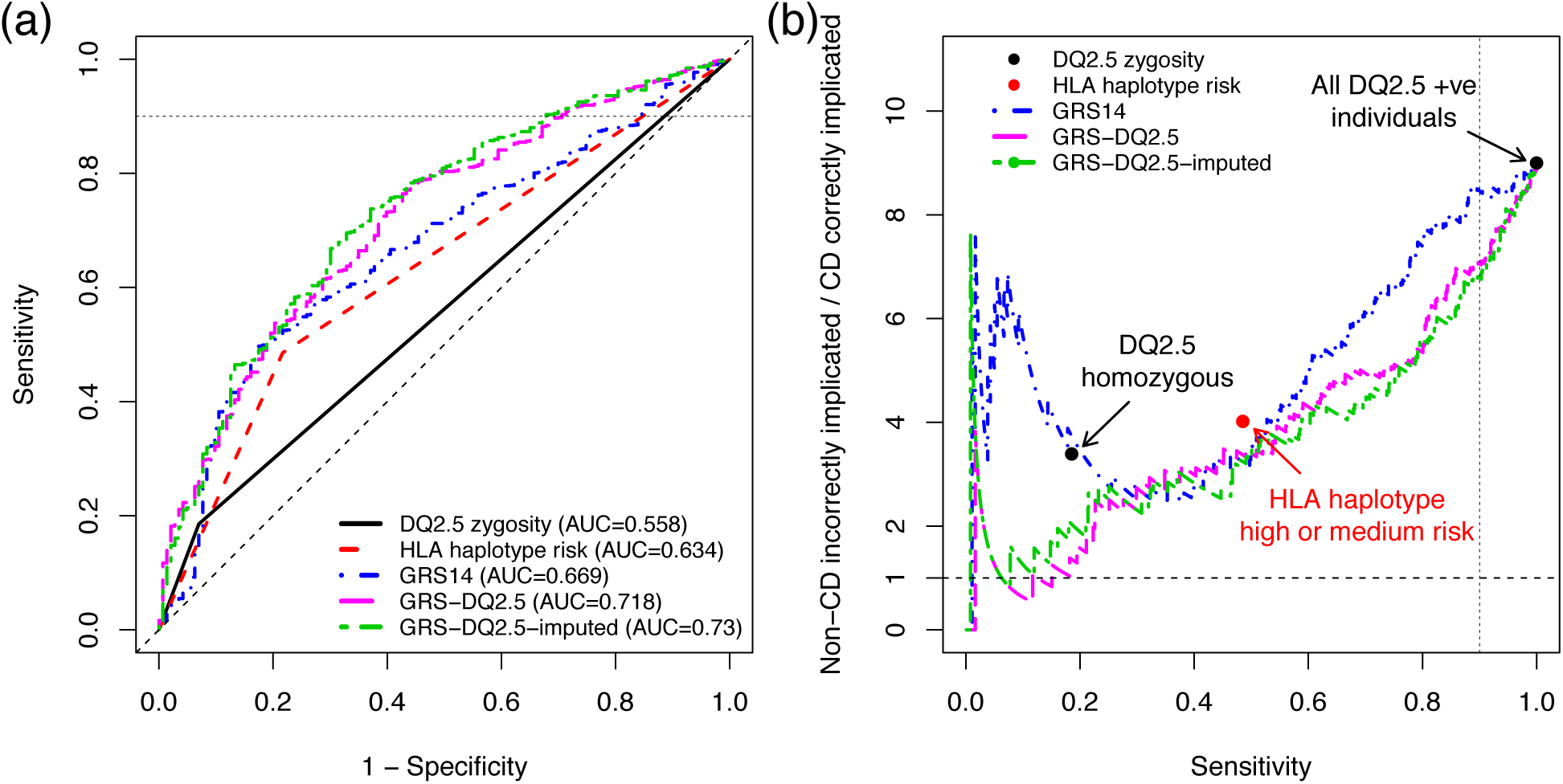
External validation results on the DQ2.5+ individuals in the North American dataset. **Legend:** (**a**) ROC curves for case/control prediction and (**b**) Non-CD implicated per CD correctly implicated, ((1 – PPV) / PPV, equivalent to 1 / [post-test-odds of disease]) versus sensitivity, for models developed on the European data and tested on the DQ2.5+ subset of the North American cohort. The DQ2.5 zygosity is the number of DQ2.5 alleles for each individual (heterozygous=1, homozygous=2). We assumed a CD prevalence of 10% in the DQ2.5+, corresponding to a baseline implication ratio of 9:1, that is, all DQ2.5+ implicated as having CD at 100% sensitivity.

In a clinical setting, it is desirable to maintain high sensitivity, that is, capturing most CD cases while incurring a cost of some false positives (reduced specificity). In considering the corresponding region of the ROC curve with sensitivity > 0.9, while both the haplotype risk model and the GRS14 model had overall slightly higher specificity than the zygosity status, the greatest increase in specificity was observed for the GRS-DQ2.5 and GRS-DQ2.5-imputed models (specificity of 0.29 and 0.32, respectively, compared with specificity = 0.15 for the GRS14).

### Utility of genomic risk scores in reducing unnecessary follow-up tests in the HLA-DQ2.5+ subgroup

In a clinical setting, a reduction in the number of unnecessary tests to screen for a disease or secure a diagnosis is desirable. The utility of a GRS for reducing unnecessary tests can be measured using the ratio of non-CD incorrectly implicated as CD to those CD cases correctly implicated (given that neither the proposed GRS nor HLA typing can act as a sole diagnostic for CD, we use the term “implicate” as distinct from “diagnosed”). This ratio should further be assessed relative to the sensitivity, as one should seek to minimize the former while maximizing the latter. A lower ratio (ideally <1) at high sensitivity indicates a better ability to avoid falsely implicating non-CD individuals as being at high CD risk, while capturing a substantial number of CD cases. Unlike the sensitivity and specificity, this ratio depends on the true prevalence of CD in the population being tested (here, all DQ2.5+ individuals). People at high-risk of CD, for instance, due to a family history of the illness, have a 10% prevalence of disease [48]. Therefore for this modeling we likewise assumed a prevalence of 10%, leading to a baseline ratio of 9:1 (equivalent to all DQ2.5-positive individuals being recommended for follow-up testing).

Overall, both the GRS-DQ2.5 and GRS-DQ2.5-imputed models implicated fewer non-CD individuals per CD case correctly implicated than the GRS14, the DQ2.5-zygosity, or the HLA-haplotype-risk score (Figure 3b). Note that since DQ2.5-zygosity within the DQ2.5+ individuals is limited to heterozygous (one risk allele) and homozygous (two risk alleles), the main point of interest for the curve is the one indicating heterozygotes, leading to ∼3.5:1 incorrect implications but achieving only 20% sensitivity. By comparison, both GRS-DQ2.5 and GRS-DQ2.5-imputed models achieved a ∼2:1 ratio at the same sensitivity as DQ2.5-zygosity. Similarly, the main point of interest for HLA-haplotype-risk separates high from medium risk, with a sensitivity of 50% and ∼4:1 incorrect implications, a level improved upon by GRS14, GRS-DQ2.5 and GRS-DQ2.5-imputed.

If we analyze the performance of the GRS at a sensitivity of 90% (comparable to that achieved by CD serology), the incorrect implications ratio was lowest for GRS-DQ2.5-imputed (6.7:1), followed by GRS-DQ2.5 (6.9:1) and GRS14 (8.5:1). To evaluate the stability of these results we performed stratified bootstrap analysis of the North American samples (*B*=10,000 replications). The greatest improvements in the average bootstrap reduction in incorrect implications ratio was from GRS-DQ2.5-imputed at a reduction of 2.17 (95% CI 1.35—3.02) over the 9:1 baseline, followed by GRS-DQ2.5 and GRS14, at 1.96 (1.16—2.78) and 1.61 (0.55—2.64) respectively (see **Additional File 1**, Supplementary Figure 7 for results limited to sensitivity ≥90%).

## Discussion

More widespread use of genomic risk prediction in autoimmune disease has been hampered by the inability to identify compelling advantages over existing approaches, mainly HLA haplotyping. Here, we have focused our analysis on individuals carrying the HLA-DQ2.5 heterodimer, which is the most common risk heterodimer and also imparts the highest risk for CD. Existing diagnostic tests are not useful in the absence of active disease and cannot predict risk of future disease. While approaches based on HLA haplotypes, including DQ2.5 zygosity and a HLA haplotype risk score [33], provide some predictive power, we have demonstrated that genomic risk scores focused on DQ2.5+ individuals have substantially higher predictive power than either approach, extending our previous findings [8]. For the DQ2.5+ subset of individuals in particular, our results highlight that while particular HLA alleles are a necessary condition for CD, they are far from a sufficient condition, and that genomic analysis can better uncover further sources of risk not adequately captured by DQ2.5 zygosity alone.

Our genomic risk scores were based on direct modeling of SNPs, both within and outside of HLA. Further, combining the genome-wide SNP data with imputed HLA markers, including HLA alleles outside the well-known *HLA-DQA1* and *HLA-DQB1* genes, imputed HLA SNPs, and HLA amino acid substitutions [35], led to an increase in predictive power on the North American dataset, thus suggesting that tools for imputing non-traditional risk factors have an important role in future predictive modeling.

The increased precision of both GRS-DQ2.5 and GRS-DQ2.5-imputed translated to an average increased saving of ∼2 unnecessary follow-ups per justified follow-up (a 22—24% average reduction), compared with the alternative strategy of considering all DQ2.5+ individuals for follow-up testing. Importantly, at this level, GRS-DQ2.5 and GRS-DQ2.5-imputed still captured 90% of DQ2.5+ CD cases. These results suggest that a GRS specific to HLA-DQ2.5+ individuals can achieve substantial cost savings while incurring only a small loss in sensitivity, relative to implicating all DQ2.5+ individuals for more intensive screening and follow-up. In real-life clinical practice, measures could be put in place to overcome the slight loss of sensitivity or any negative consequences. For instance, combining the GRS with CD serology would enable the high sensitivity of the latter test to minimize the chance of true CD cases being overlooked but still leverage the discriminatory power of the GRS. Patients would still be advised to seek medical review if they developed symptoms suggestive of CD. Further, the top 15—20% of CD cases in terms of GRS-DQ2.5 were estimated to be detectable at a level of 1 unnecessary follow-up per true CD case, suggesting that relatively high confidence of CD can be conferred upon individuals with the highest GRS-DQ2.5/GRS-DQ2.5-imputed scores, thus potentially only requiring CD serology for the subset of individuals with moderate risk, who cannot be confidently predicted by the GRS to have CD or be healthy.

The ultimate clinical role, utility, and cost-effectiveness of genomic risk scores for CD remain to be determined in prospective clinical studies where genomic profiles are undertaken from the outset. If prospective studies can establish the clinical value of the GRS and link it to CD risk, we envisage that clinicians ordering this test will be provided with a weighted GRS value that links to a scale of CD risk. They would then use this score to complement clinical risk stratification and guide the most suitable advice on long-term follow-up and/or further investigations.

Several potential roles for the GRS can be proposed. First, guiding ongoing care in people at-risk of CD who are also HLA-DQ2.5+. Several recent large prospective studies have reinforced the importance of HLA-DQ2.5 for CD development in childhood, with environmental factors such as age of gluten introduction or breastfeeding having little, if any, impact on the proportion of children who ultimately develop CD [25, 26, 49]. These findings have prompted some experts to recommend increased surveillance for CD in children with a positive family history of CD who are HLA-DQ2.5, especially if DQ2.5 homozygous. The increased predictive power of our GRS-DQ2.5 beyond the information afforded by DQ2.5 zygosity status alone is likely to provide greater clinical utility for predicting risk in this important subgroup of individuals.

Second, genomic data may also be able to inform CD prognosis, such as the likelihood of complicated (refractory) disease or the natural history of latent CD (positive CD-serology and HLA susceptibility but a normal small bowel), or other aspects of care such as response to the gluten-free diet. Finally, another approach may be to combine a GRS with serology to optimize risk stratification for CD and determine who will benefit most from definitive small bowel biopsy. Such a strategy could leverage the strengths of each test: the high sensitivity for active CD using CD serology and the fine-grained CD risk quantification of the GRS, including its ability to provide predictive information and exclude CD. A major benefit of genetic testing in the diagnostic work-up of CD is that, unlike serology and small bowel histology, accurate results are not dependent on active gluten intake. This is particularly relevant, as the gluten free diet has been adopted by 10% or more of the population in many Western countries, rendering traditional tests inaccurate. Establishing the clinical utility of genomic testing in CD will also support the feasibility of a genomics-based platform for a range of other autoimmune diseases, where both HLA and non-HLA genetic contributions are important as well, and which commonly overlap with CD [14].

## Conclusions

Our findings highlight the value of genomic risk scores that target a clinically relevant subgroup of individuals at-risk for CD. Genomic risk scores that utilize genome-wide SNPs, HLA alleles, and amino acid substitutions provided the highest predictive power in individuals who are DQ2.5+, surpassing that of approaches based on small numbers of well-known risk haplotypes or models of SNPs only. This improved predictive power directly translates to an ability to better stratify DQ2.5+ individuals by CD risk, meaning that for each justified test, two follow-up tests in people unlikely to develop CD could be avoided, which improves both patient care and health care delivery. Future clinical studies will enable optimization of such risk scores to particular clinical settings and assess how to best integrate genomic risk prediction with the current clinical diagnostic pathways for CD. Our results in CD suggest that employing such genomic-based approaches in other autoimmune disease is both feasible and potentially of clinical utility.

## Author contributions

Designed the study: GA, JT-D, MI. Performed experiments and analyzed data: GA, AR. All authors contributed to and approved the final manuscript.

## Additional files

**Additional file 1**: Supplementary results and figures.

**Additional file 2:** The GRS-DQ2.5 score, in terms of SNP ID, reference allele, and weight.

**Additional file 3:** The GRS-DQ2.5-imputed score, in terms of SNP/marker ID, reference allele, and weight.

## Acknowledgments

The North American Celiac Disease Consortium study was conducted by the North American Celiac Disease Consortium Investigators and supported by the National Institute of Diabetes and Digestive and Kidney Diseases (NIDDK). The data from the North American Celiac Disease Consortium reported here were supplied by the NIDDK Central Repositories. This manuscript was not prepared in collaboration with Investigators of the North American Celiac Disease Consortium study and does not necessarily reflect the opinions or views of the North American Celiac Disease Consortium study, the NIDDK Central Repositories, or the NIDDK.

We thank the chief investigators of the van Heel et al., 2007, Dubois et al., 2010 and Trynka et al., 2011 papers (David van Heel, Cisca Wijmenga, and Lude Franke) for providing the celiac disease data.

The authors acknowledge support and funding from NHMRC grant no. 1062227. GA was supported by an NHMRC Early Career Fellowship (no. 1090462). MI was supported by a Career Development Fellowship co-funded by the NHMRC and Heart Foundation (no. 1061435).

## Supplementary Results

### Assessment of potential reasons for the reduction in AUC on the North American dataset

We sought to investigate the source for the small reduction in AUC observed in the North American dataset relative to the European data. First, we employed the same methods used to develop the GRS14 (L1-penalized linear support vector machines) to generate a CD risk score within the North American dataset; specifically, we used 10-fold cross-validation within the North American dataset to estimate the predictive power (area under receiver-operating-characteristic curve, AUC) of the penalized models, as a function of the number of SNPs assigned a non-zero weight in the model (Supplementary Figure 3). The best average AUC of 0.823 was achieved with models including ∼40 SNPs with non-zero weights. Beyond that, increasing the number of SNPs with non-zero weight in the model reduced the cross-validated AUC, indicating overfitting. As further verification that the difference in AUC was not driven by our choice of model, we employed MultiBLUP, a non-sparse genome-wide modeling method based on linear mixed models, on the European GWA data (n=11,912), and tested its predictions on the North American cohort. MultiBLUP achieved AUC = 0.831 (95% CI 0.808—0.853), equal to the AUC of the GRS14. Next, we examined the distribution of the GRS14 scores, stratified by CD diagnosis method (controls, biopsy & serology, biopsy, serology, and unknown), to examine whether there were substantial differences in predicted risk between diagnosis methods that could indicate potential misclassification of case/control due to variability in diagnosis method. We did not observe substantial differences in the median risk between diagnosis methods apart from the expected difference between cases (regardless of diagnosis method) and controls (Supplementary Figure 4). Finally, we computed the fixation index F_st_ for the 224 SNPs in the risk score between the North American and European data (mean F_st_=3×10^−3^ based on 213 SNPs with valid estimates), confirming the earlier PCA results showing negligible allele frequency differences between the training and validation datasets.

### GRS-DQ2.5 based on a combined Immunochip and GWA training set

We also explored whether using a DQ2.5-specific GRS trained on the combined European GWA and Immunochip datasets, employing the SNPs common to the two platforms and other SNP2HLA markers (10,284 samples and 24,555 SNPs+markers), would lead to increased predictive power over using just the European GWA data. While training SparSNP on this combined SNP+marker dataset led to increases in cross-validated AUC (0.748 for an L2 penalty of 0.01; Supplementary Figure 6), this model subsequently achieved lower AUC in external validation on the North American dataset (AUC = 0.707, 95% CI 0.663—0.750).

**Supplementary Figure 1:**
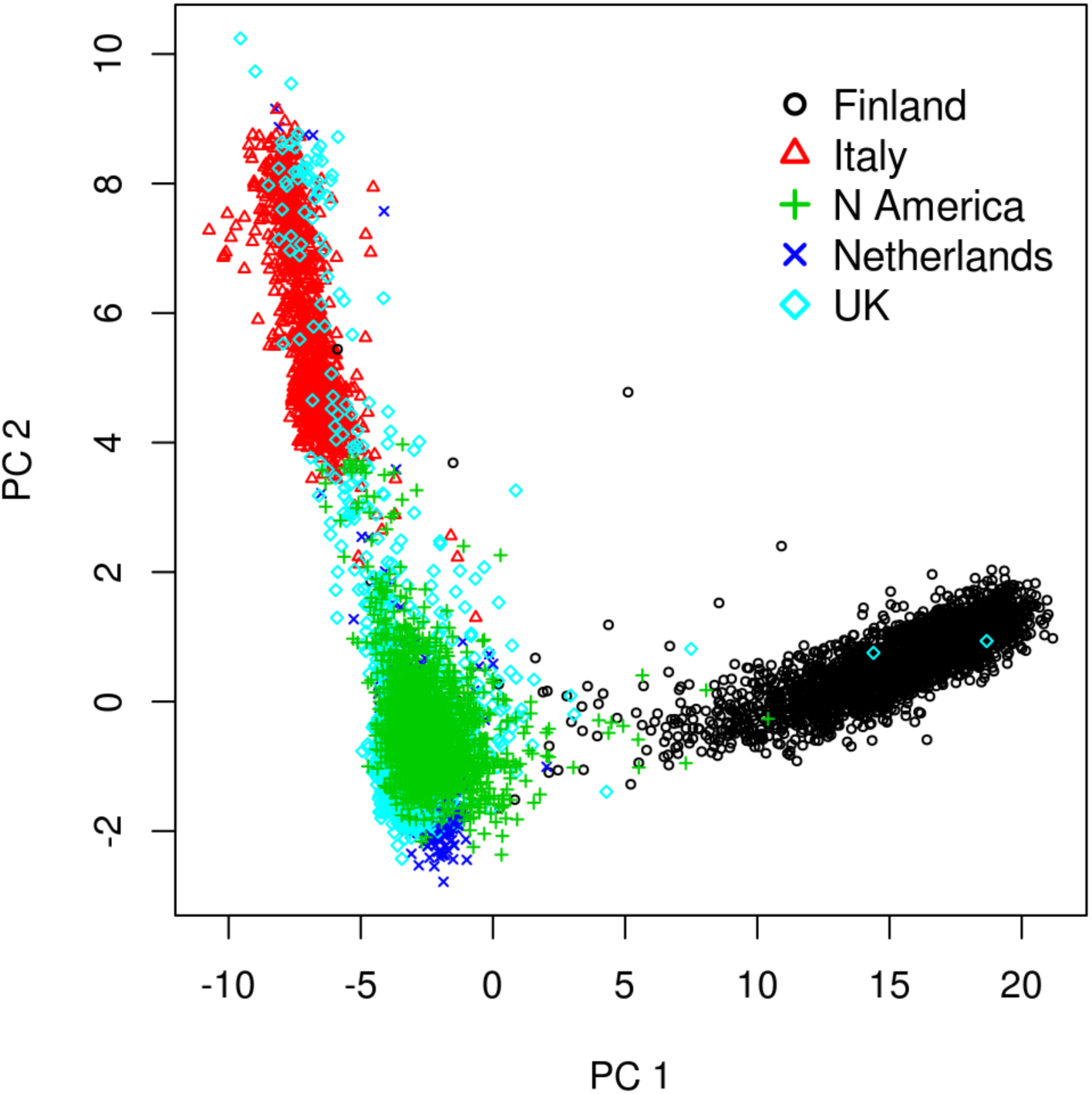
The first two principal components (PCs) of an LD-thinned dataset combining the European GWA datasets (Finland, Italy, Netherlands, and UK) and the North American dataset (post QC).

**Supplementary Figure 2:**
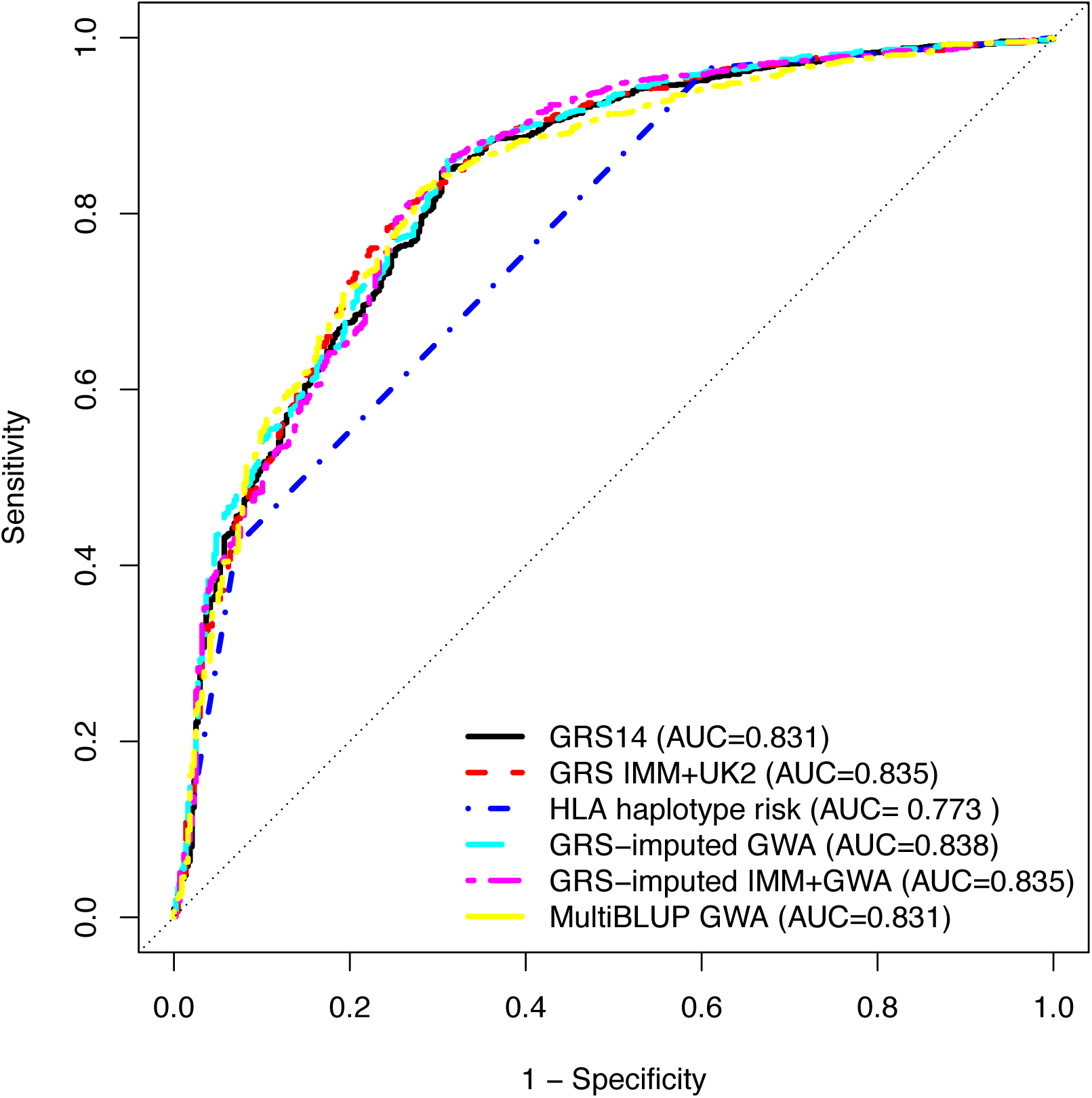
ROC curves for classifying all CD cases and controls using different predictors in the North American dataset. *GRS14*: the published GRS (trained on the UK2 dataset); *GRS IMM+UK2*: a GRS trained on the Immunochip + UK SNP data; *GRS-imputed GWA:* a GRS trained on all European GWA datasets (UK, Dutch, Finnish, Italian), consisting of SNPs and SNP2HLA imputed markers; *GRS-imputed IMM+GWA*: a GRS trained on the European GWA + Immunochip datasets, consisting of SNPs and SNP2HLA imputed markers; *HLA haplotype risk:* a 3-level risk score based on the imputed HLA haplotype status; *MultiBLUP GWA*: a MultiBLUP model trained on the European GWA data.

**Supplementary Figure 3:**
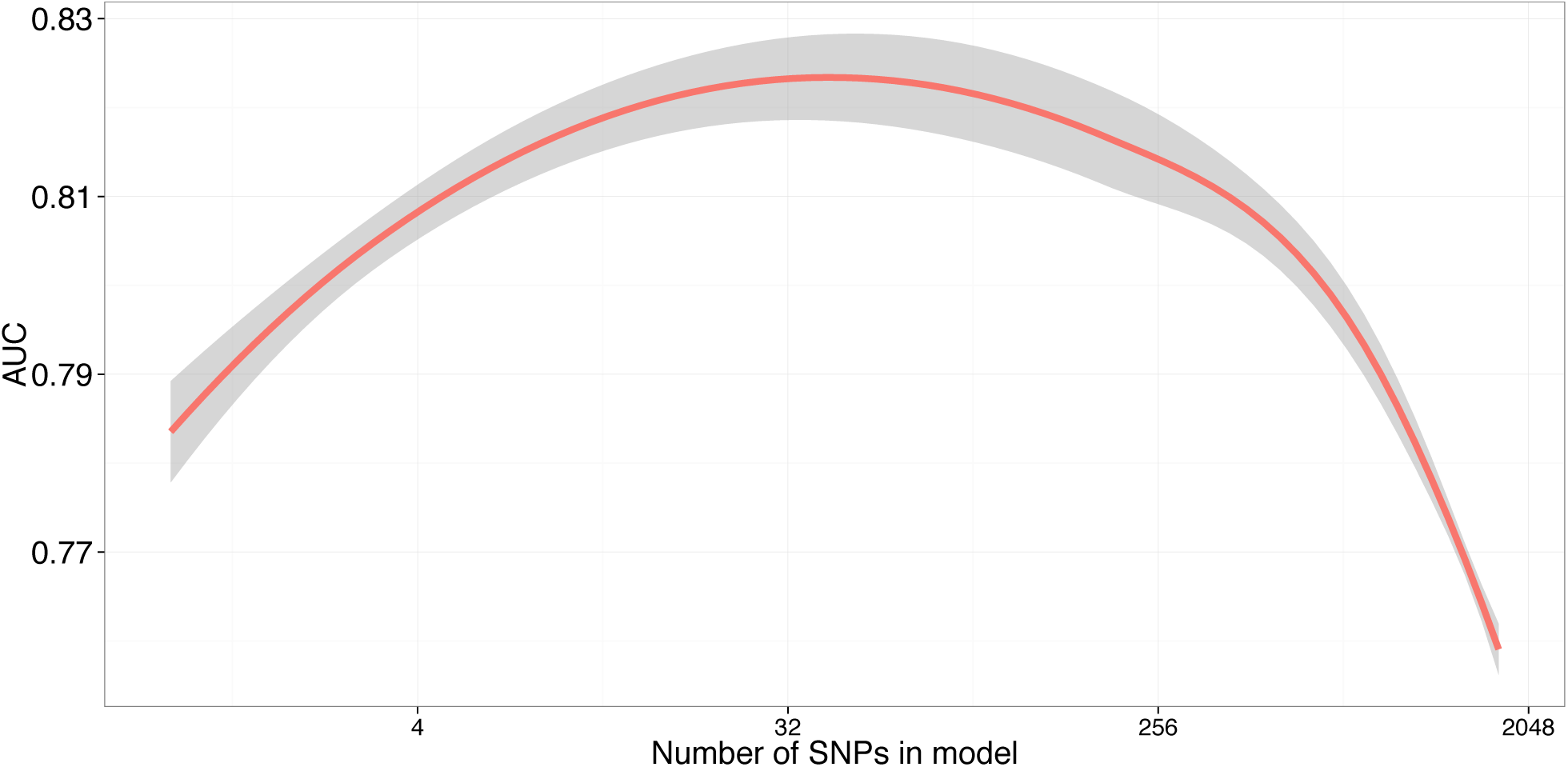
10×10 cross-validated AUC (LOESS-smoothed) for the GRS developed within the North American dataset, as a function of the number of SNPs assigned a non-zero weight in the model. The maximum AUC was 0.823 at 40 SNPs with non-zero weight.

**Supplementary Figure 4:**
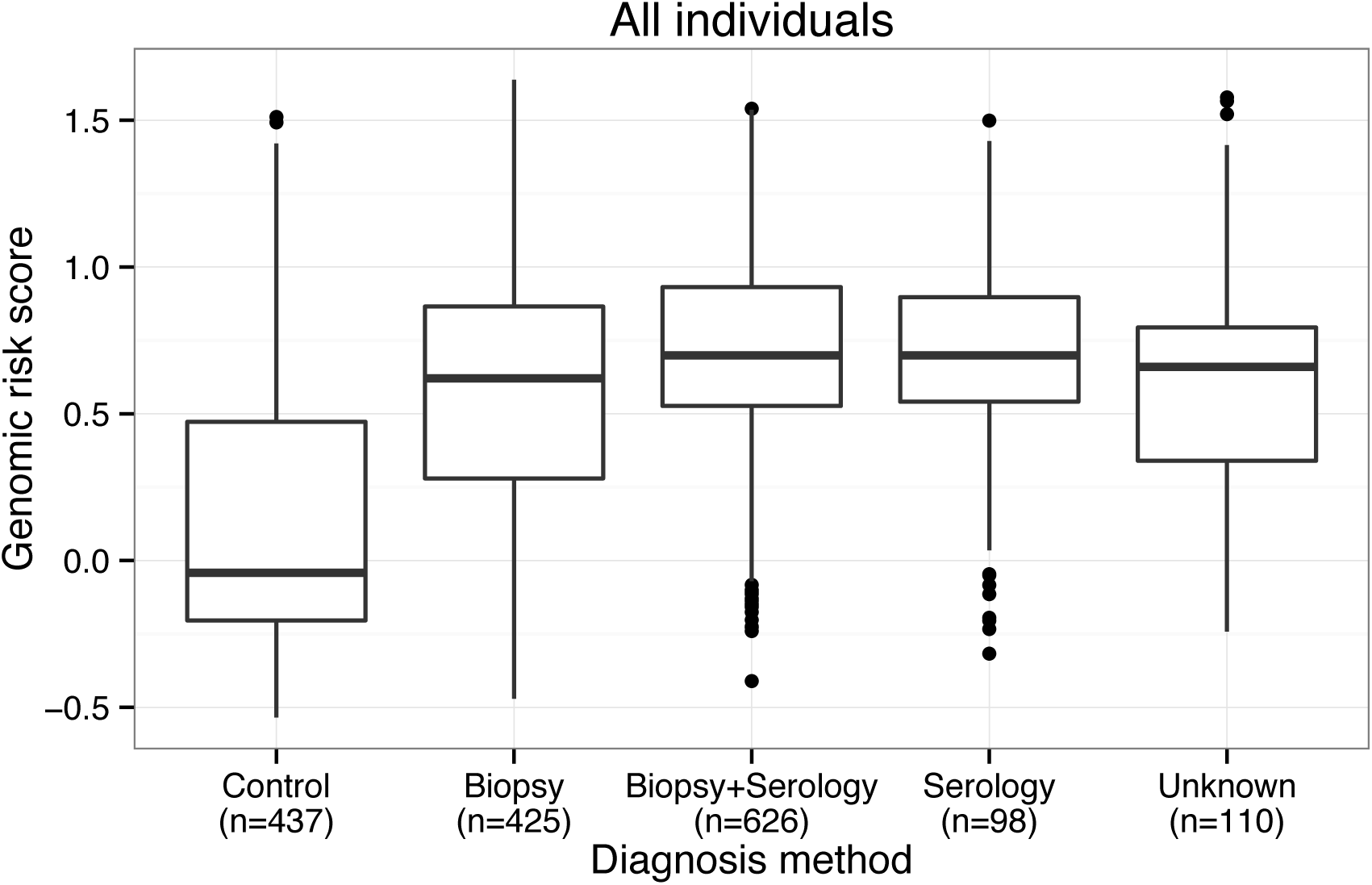
Boxplots of the genomic risk scores (GRS14) within each diagnosis method for all individuals in the North American cohort (n=1696).

**Supplementary Figure 5:**
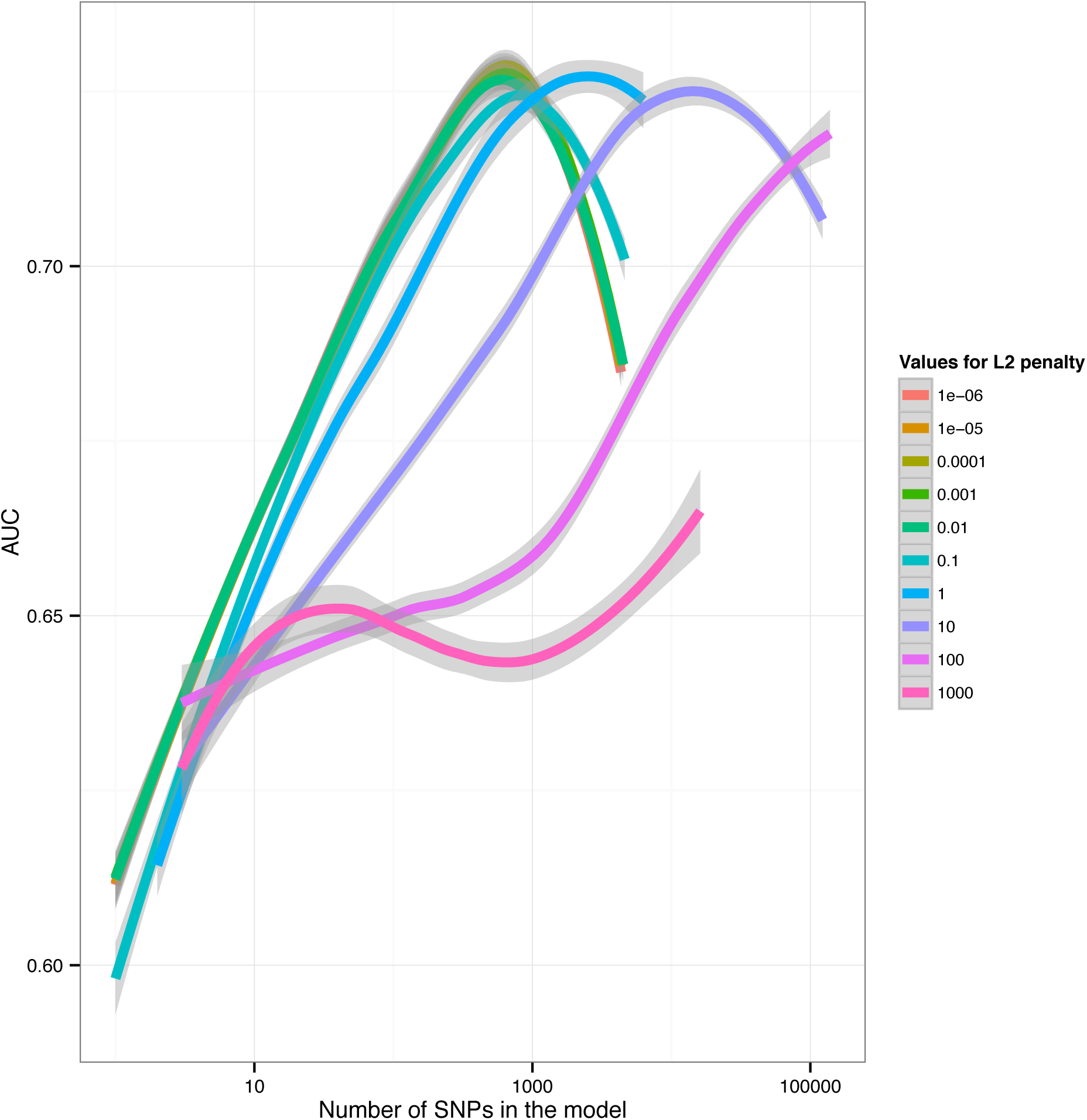
10×10 cross-validated AUC (LOESS-smoothed) for 10 candidate GRS developed on the DQ-2.5+ individuals from the European GWA SNP datasets (n=5552), each using a different L2 penalty.

**Supplementary Figure 6:**
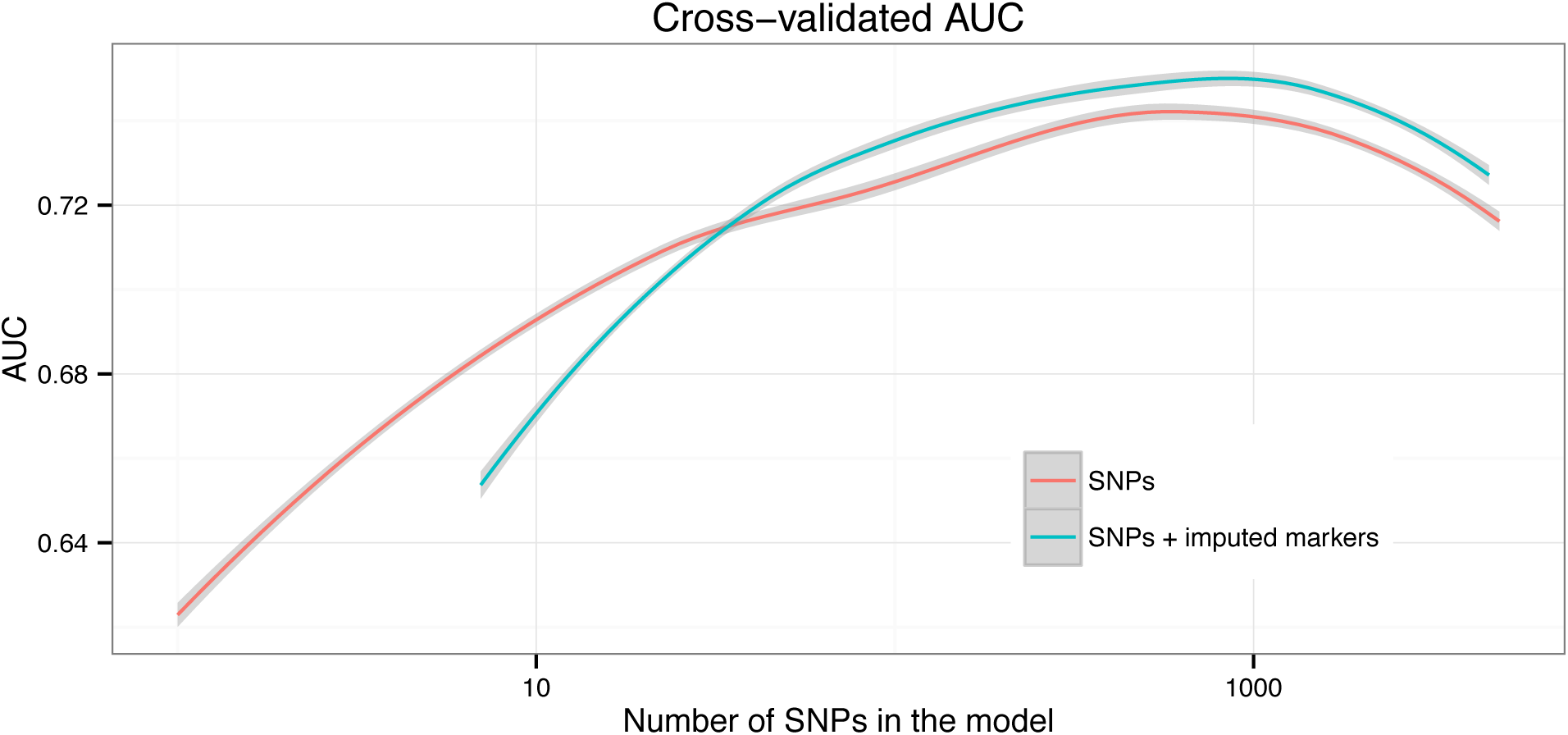
10×10 cross-validated AUC (LOESS-smoothed) for the novel GRS-DQ2.5 model trained on the combined DQ2.5+ subsets of the European GWA data and the Immunochip data (n=10,284), as a function of the number of SNPs assigned a non-zero weight in the model. For the SNP+marker model, maximum AUC of 0.742 was achieved at 583 SNPs/markers with non-zero weight using an L2 penalty of 0.01.

**Supplementary Figure 7:**
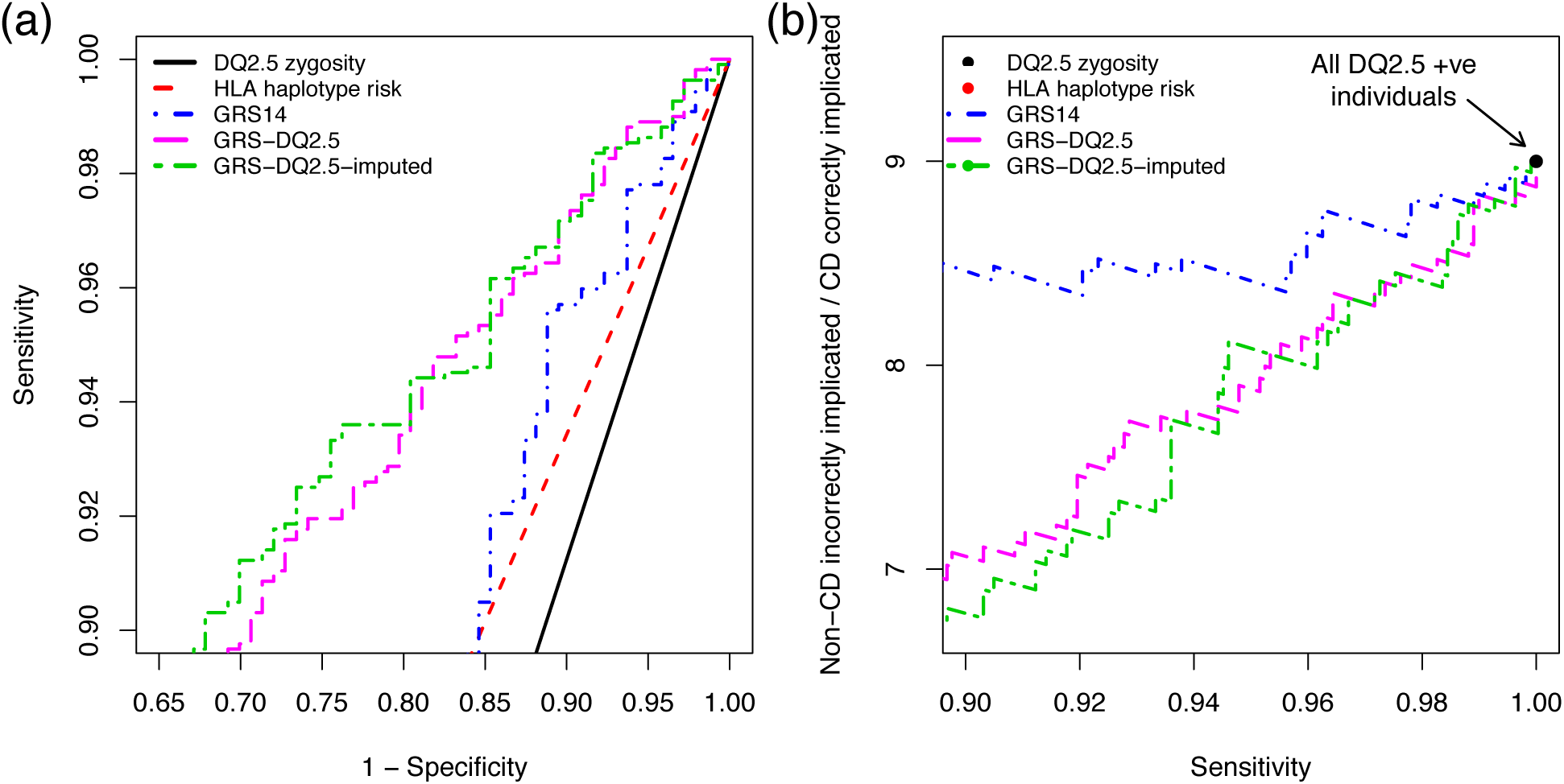
External validation results on the DQ2.5+ individuals in the North American dataset, focusing on sensitivity ≥90%. (**a**) ROC curves for case/control prediction and (**b**) Non-CD implicated per CD correctly implicated, ((1 – PPV) / PPV, equivalent to 1 / [post-test-odds of disease]) versus sensitivity, for models developed on the European data and tested on the DQ2.5+ subset of the North American cohort. The DQ2.5 zygosity is the number of DQ2.5 alleles for each individual (heterozygous=1, homozygous=2). We assumed a CD prevalence of 10% in the DQ2.5+, corresponding to a baseline implication ratio of 9:1, that is, all DQ2.5+ implicated as having CD at 100% sensitivity. Note that the estimate of HLA haplotype risk for (b) fall below the sensitivity of 90% and are not shown.

## References

1. Graham, R.R., C. Cotsapas, L. Davies, R. Hackett, C.J. Lessard, J.M. Leon, N.P. Burtt, C. Guiducci, et al., Genetic variants near TNFAIP3 on 6q23 are associated with systemic lupus erythematosus. Nat Genet, 2008. 40(9): p. 1059–61.

2. Ramos, P.S., L.A. Criswell, K.L. Moser, M.E. Comeau, A.H. Williams, N.M. Pajewski,S.A. Chung, R.R. Graham, et al., A comprehensive analysis of shared loci between systemic lupus erythematosus (SLE) and sixteen autoimmune diseases reveals limited genetic overlap. PLoS Genetics, 2011. 7(12): p. e1002406.

3. Trynka, G., K.a. Hunt, N.a. Bockett, J. Romanos, V. Mistry, A. Szperl, S.F. Bakker, M.T. Bardella, et al., Dense genotyping identifies and localizes multiple common and rare variant association signals in celiac disease. Nature Genetics, 2011. 43: p. 1193–201.

4. Dubois, P.C.A., G. Trynka, L. Franke, K.a. Hunt, J. Romanos, A. Curtotti, A. Zhernakova,G.a.R. Heap, et al., Multiple common variants for celiac disease influencing immune gene expression. Nature Genetics, 2010. 42: p. 295–302.

5. Barrett, J.C., D.G. Clayton, P. Concannon, B. Akolkar, J.D. Cooper, H.A. Erlich, C. Julier,G. Morahan, et al., Genome-wide association study and meta-analysis find that over 40 loci affect risk of type 1 diabetes. Nature Genetics, 2009. 41(6): p. 703–7.

6. Zhernakova, A., E.A. Stahl, G. Trynka, S. Raychaudhuri, E.A. Festen, L. Franke, H.J. Westra, R.S. Fehrmann, et al., Meta-analysis of genome-wide association studies in celiac disease and rheumatoid arthritis identifies fourteen non-HLA shared loci. PLoS Genetics, 2011. 7(2): p. e1002004.

7. Wei, Z., K. Wang, H.Q. Qu, H. Zhang, J. Bradfield, C. Kim, E. Frackleton, C. Hou, et al., From disease association to risk assessment: an optimistic view from genome-wide association studies on type 1 diabetes. PLoS Genet, 2009. 5(10): p. e1000678.

8. Abraham, G., J.A. Tye-Din, O.G. Bhalala, A. Kowalczyk, J. Zobel, and M. Inouye, Accurate and robust genomic prediction of celiac disease using statistical learning. PLoS Genet, 2014. 10(2): p. e1004137.

9. Wei, Z., W. Wang, J. Bradfield, J. Li, C. Cardinale, E. Frackelton, C. Kim, F. Mentch, et al., Large sample size, wide variant spectrum, and advanced machine-learning technique boost risk prediction for inflammatory bowel disease. Am J Hum Genet, 2013. 92(6): p. 1008–12.

10. Hoggart, C.J., J.C. Whittaker, M. De Iorio, and D.J. Balding, Simultaneous analysis of all SNPs in genome-wide and re-sequencing association studies. PLoS Genetics, 2008. 4(7): p. e1000130.

11. Speed, D. and D.J. Balding, MultiBLUP: improved SNP-based prediction for complex traits. Genome Res, 2014. 24(9): p. 1550–7.

12. Abraham, G., A. Kowalczyk, J. Zobel, and M. Inouye, Performance and robustness of penalized and unpenalized methods for genetic prediction of complex human disease. Genetic Epidemiology, 2013. 37(2): p. 184–195.

13. Husby, S., S. Koletzko, I.R. Korponay-Szabo, M.L. Mearin, A. Phillips, R. Shamir, R. Troncone, K. Giersiepen, et al., European Society for Pediatric Gastroenterology, Hepatology, and Nutrition guidelines for the diagnosis of coeliac disease. J Pediatr Gastroenterol Nutr, 2012. 54(1): p. 136–60.

14. Abadie, V., L.M. Sollid, L.B. Barreiro, and B. Jabri, Integration of genetic and immunological insights into a model of celiac disease pathogenesis. Annu Rev Immunol, 2011. 29: p. 493–525.

15. van Heel, D.a. and J. West, Recent advances in coeliac disease. Gut, 2006. 55: p. 1037–46.

16. Nistico, L., C. Fagnani, I. Coto, S. Percopo, R. Cotichini, M.G. Limongelli, F. Paparo, S. D’Alfonso, et al., Concordance, disease progression, and heritability of coeliac disease in Italian twins. Gut, 2006. 55(6): p. 803–8.

17. Karell, K., A.S. Louka, S.J. Moodie, H. Ascher, F. Clot, L. Greco, P.J. Ciclitira, L.M. Sollid, et al., HLA types in celiac disease patients not carrying the DQA1*05-DQB1*02 (DQ2)heterodimer: results from the European Genetics Cluster on Celiac Disease. Hum Immunol, 2003. 64(4): p. 469–77.

18. Tye-Din, J.A., J.A. Stewart, J.A. Dromey, T. Beissbarth, D.A. van Heel, A. Tatham, K. Henderson, S.I. Mannering, et al., Comprehensive, quantitative mapping of T cell epitopes in gluten in celiac disease. Sci Transl Med, 2010. 2(41): p. 41ra51.

19. Hunt, K.A., A. Zhernakova, G. Turner, G.A. Heap, L. Franke, M. Bruinenberg, J. Romanos,L.C. Dinesen, et al., Newly identified genetic risk variants for celiac disease related to the immune response. Nat Genet, 2008. 40(4): p. 395–402.

20. Ahn, R., Y.C. Ding, J. Murray, A. Fasano, P.H. Green, S.L. Neuhausen, and C. Garner, Association analysis of the extended MHC region in celiac disease implicates multiple independent susceptibility loci. PLoS One, 2012. 7(5): p. e36926.

21. Festen, E.A., P. Goyette, T. Green, G. Boucher, C. Beauchamp, G. Trynka, P.C. Dubois, C. Lagace, et al., A meta-analysis of genome-wide association scans identifies IL18RAP, PTPN2, TAGAP, and PUS10 as shared risk loci for Crohn’s disease and celiac disease. PLoS Genet, 2011. 7(1): p. e1001283.

22. Garner, C., R. Ahn, Y.C. Ding, L. Steele, S. Stoven, P.H. Green, A. Fasano, J.A. Murray, et al., Genome-wide association study of celiac disease in North America confirms FRMD4B as new celiac locus. PLoS One, 2014. 9(7): p. e101428.

23. Tye-Din, J.A., D.J. Cameron, A.J. Daveson, A.S. Day, P. Dellsperger, C. Hogan, E.D. Newnham, S.J. Shepherd, et al., Appropriate clinical use of human leukocyte antigen typing for coeliac disease: an Australasian perspective. Intern Med J, 2015. 45(4): p. 441–50.

24. Anderson, R.P., M.J. Henry, R. Taylor, E.L. Duncan, P. Danoy, M.J. Costa, K. Addison,J.A. Tye-Din, et al., A novel serogenetic approach determines the community prevalence of celiac disease and informs improved diagnostic pathways. BMC Med, 2013. 11(1): p. 188.

25. Lionetti, E., S. Castellaneta, R. Francavilla, A. Pulvirenti, E. Tonutti, S. Amarri, M. Barbato,C. Barbera, et al., Introduction of gluten, HLA status, and the risk of celiac disease in children. N Engl J Med, 2014. 371(14): p. 1295–303.

26. Liu, E., H.S. Lee, C.A. Aronsson, W.A. Hagopian, S. Koletzko, M.J. Rewers, G.S. Eisenbarth, P.J. Bingley, et al., Risk of pediatric celiac disease according to HLA haplotype and country. N Engl J Med, 2014. 371(1): p. 42–9.

27. Karinen, H., P. Karkkainen, J. Pihlajamaki, E. Janatuinen, M. Heikkinen, R. Julkunen, V.M. Kosma, A. Naukkarinen, et al., Gene dose effect of the DQB1*0201 allele contributes to severity of coeliac disease. Scand J Gastroenterol, 2006. 41(2): p. 191–9.

28. Al-Toma, A., M.S. Goerres, J.W. Meijer, A.S. Pena, J.B. Crusius, and C.J. Mulder, Human leukocyte antigen-DQ2 homozygosity and the development of refractory celiac disease and enteropathy-associated T-cell lymphoma. Clin Gastroenterol Hepatol, 2006. 4(3): p. 315–9.

29. Biagi, F., A. Schiepatti, G. Malamut, A. Marchese, C. Cellier, S.F. Bakker, C.J. Mulder, U. Volta, et al., PROgnosticating COeliac patieNts SUrvivaL: the PROCONSUL score. PLoS One, 2014. 9

30. Abraham, G., A. Kowalczyk, J. Zobel, and M. Inouye, SparSNP: Fast and memory-efficient analysis of all SNPs for phenotype prediction. BMC Bioinformatics, 2012. 13: p. 88.

31. Okser, S., T. Pahikkala, A. Airola, T. Salakoski, S. Ripatti, and T. Aittokallio, Regularized Machine Learning in the Genetic Prediction of Complex Traits. PLoS Genet, 2014. 10(11): p. e1004754.

32. van Heel, D.A., L. Franke, K.A. Hunt, R. Gwilliam, A. Zhernakova, M. Inouye, M.C. Wapenaar, M.C.N.M. Barnardo, et al., A genome-wide association study for celiac disease identifies risk variants in the region harboring IL2 and IL21. Nature Genetics, 2007. 39: p. 827–9.

33. Romanos, J., A. Rosen, V. Kumar, G. Trynka, L. Franke, A. Szperl, J. Gutierrez-Achury,C.C. van Diemen, et al., Improving coeliac disease risk prediction by testing non-HLA variants additional to HLA variants. Gut, 2014. 63(3): p. 415–22.

34. Gutierrez-Achury, J., A. Zhernakova, S.L. Pulit, G. Trynka, K.A. Hunt, J. Romanos, S. Raychaudhuri, D.A. van Heel, et al., Fine mapping in the MHC region accounts for 18% additional genetic risk for celiac disease. Nat Genet, 2015. 47(6): p. 577–8.

35. Jia, X., B. Han, S. Onengut-Gumuscu, W.M. Chen, P.J. Concannon, S.S. Rich, S. Raychaudhuri, and P.I. de Bakker, Imputing amino acid polymorphisms in human leukocyte antigens. PLoS One, 2013. 8(6): p. e64683.

36. Garner, C.P., J.A. Murray, Y.C. Ding, Z. Tien, D.A. van Heel, and S.L. Neuhausen, Replication of celiac disease UK genome-wide association study results in a US population. Hum Mol Genet, 2009. 18(21): p. 4219–25.

37. Wray, N.R., J. Yang, B.J. Hayes, A.L. Price, M.E. Goddard, and P.M. Visscher, Pitfalls of predicting complex traits from SNPs. Nat Rev Genet, 2013. 14(7): p. 507–15.

38. Purcell, S., B. Neale, K. Todd-Brown, L. Thomas, M.A. Ferreira, D. Bender, J. Maller, P. Sklar, et al., PLINK: a tool set for whole-genome association and population-based linkage analyses. Am J Hum Genet, 2007. 81(3): p. 559–75.

39. Chang, C.C., C.C. Chow, L.C. Tellier, S. Vattikuti, S.M. Purcell, and J.J. Lee, Second-generation PLINK: rising to the challenge of larger and richer datasets. Gigascience, 2015. 4: p. 7.

40. Abraham, G. and M. Inouye, Fast principal component analysis of large-scale genome-wide data. PLoS One, 2014. 9(4): p. e93766.

41. Weir, B.S. and C.C. Cockerham, Estimating F-Statistics for the Analysis of Population-Structure. Evolution, 1984. 38(6): p. 1358–1370.

42. Zheng, X., J. Shen, C. Cox, J.C. Wakefield, M.G. Ehm, M.R. Nelson, and B.S. Weir, HIBAG-HLA genotype imputation with attribute bagging. Pharmacogenomics J, 2014. 14(2): p. 192–200.

43. Harrell, F.E.J., Regression Modeling Strategies. Springer Series in Statistics, 2001: Springer.

44. Robin, X., N. Turck, A. Hainard, N. Tiberti, F. Lisacek, J.C. Sanchez, and M. Muller, pROC: an open-source package for R and S+ to analyze and compare ROC curves. BMC Bioinformatics, 2011. 12: p. 77.

45. Megiorni, F., B. Mora, M. Bonamico, M. Barbato, R. Nenna, G. Maiella, P. Lulli, and M.C. Mazzilli, HLA-DQ and risk gradient for celiac disease. Human immunology, 2009. 70: p. 55–9.

46. Fasano, A., I. Berti, T. Gerarduzzi, T. Not, R.B. Colletti, S. Drago, Y. Elitsur, P.H. Green, et al., Prevalence of celiac disease in at-risk and not-at-risk groups in the United States: a large multicenter study. Arch Intern Med, 2003. 163(3): p. 286–92.

47. Husby, S. and J.A. Murray, Diagnosing coeliac disease and the potential for serological markers. Nat Rev Gastroenterol Hepatol, 2014. 11(11): p. 655–663.

48. Sollid, L.M. and B.A. Lie, Celiac disease genetics: current concepts and practical applications. Clin Gastroenterol Hepatol, 2005. 3(9): p. 843–51.

49. Vriezinga, S.L., R. Auricchio, E. Bravi, G. Castillejo, A. Chmielewska, P. Crespo Escobar,S. Kolacek, S. Koletzko, et al., Randomized feeding intervention in infants at high risk for celiac disease. N Engl J Med, 2014. 371(14): p. 1304–15.

